# Hierarchical transcriptional regulation of quorum-sensing genes in *Vibrio harveyi*

**DOI:** 10.1101/2020.01.22.916312

**Authors:** Ryan R. Chaparian, Alyssa Ball, Julia C. van Kessel

**Affiliations:** Department of Biology, Indiana University, Bloomington, IN

**Keywords:** quorum sensing, gene regulation, LuxR, Vibrio harveyi

## Abstract

In vibrios, quorum sensing controls hundreds of genes that are required for cell density-specific behaviors including bioluminescence, biofilm formation, competence, secretion, and swarming motility. The central transcription factor in the quorum-sensing pathway is LuxR/HapR, which directly regulates ∼100 genes in the >400-gene regulon of *Vibrio harveyi*. Among these directly controlled genes are 15 transcription factors, which we predicted would comprise the second tier in the hierarchy of the quorum-sensing regulon. We confirmed that LuxR binds to the promoters of these genes *in vitro* and quantified the extent of LuxR activation or repression of transcript levels. RNA-seq indicates that most of these transcriptional regulators control only a few genes, with the exception of MetJ, which is a global regulator. The genes regulated by these transcription factors are predicted to be involved in methionine and thiamine biosynthesis, membrane stability, RNA processing, c-di-GMP degradation, sugar transport, and other cellular processes. These data support a hierarchical model in which LuxR directly regulates 15 transcription factors that drive the second level of the gene expression cascade to influence cell density-dependent metabolic states and behaviors in *V. harveyi*.

**Importance:** Quorum sensing is important for survival of bacteria in nature and influences the actions of bacterial groups. In the relatively few studied examples of quorum sensing-controlled genes, these genes are associated with competition or cooperation in complex microbial communities and/or virulence in a host. However, quorum sensing in vibrios controls the expression of hundreds of genes, and their functions are mostly unknown or uncharacterized. In this study, we identify the regulators of the second-tier of gene expression in the quorum-sensing system of the aquatic pathogen *Vibrio harveyi.* Our identification of regulatory networks and metabolic pathways controlled by quorum sensing can be extended and compared to other *Vibrio* species to understand the physiology, ecology, and pathogenesis of these organisms.

## Introduction

Bacteria coordinate gene expression in response to an array of external stimuli including nutrients, pH, and temperature. Other environmental cues that control gene expression are autoinducers (AIs) – signaling molecules that are produced by bacterial communication systems termed quorum sensing (QS) (1). In nature, bacteria live in complex microbial communities where population-wide synchronization is critical for survival. While QS circuitry differs between organisms, the core function remains constant – to enable locally unified gene expression. The QS system of *Vibrio harveyi*, a significant marine pathogen, has been studied for decades and has yielded a wealth of knowledge about communication in Gram-negative bacteria (2–4). In this bacterium, three distinct AIs are produced intracellularly and diffuse into the local surroundings. These three AIs each bind cognate membrane-bound histidine sensor kinase receptors located in the inner membrane. The receptor proteins possess dual functionality as kinases and phosphatases, which controls the flow of phosphate through the quorum sensing circuit (5). At low cell density (LCD), AI concentration is low and receptor proteins remain predominantly unbound by AIs. In the unbound state, these proteins function as kinases which phosphorylate LuxU, a phosphotransfer protein, which is responsible for subsequently phosphorylating LuxO. In its phosphorylated state, LuxO, a response regulator, activates transcription of the five quorum regulatory RNAs (Qrrs) (6). The Qrrs stabilize *aphA* mRNA and destabilize *luxR* mRNA (7). The end result is high level production of AphA at LCD, the LCD master regulator, and simultaneous suppression of LuxR, the high cell density (HCD) master regulator. At LCD, cells configure gene expression programs via AphA and low levels of LuxR to behave as individuals (3). Conversely, as bacterial populations mature, AI concentrations exceed threshold concentrations in which cognate receptor proteins are highly bound by AIs. AI binding promotes the phosphatase activities of the receptor proteins (8). Under these conditions, LuxO is dephosphorylated, which ultimately results in cessation of AphA production and derepression of *luxR*. This derepression allows for a 10-fold change in *luxR* expression which translates to a spectrum of protein ranging between 600 and 6,000 dimers per cell (9, 10). Once at quorum, LuxR is maximally expressed and gene expression programs are adjusted to produce group behaviors. This circuitry is conserved across vibrios, and LuxR/HapR-type proteins are the core regulators of genes at HCD in all vibrios, including the human pathogens *Vibrio cholerae* (HapR), *Vibrio parahaemolyticus* (OpaR), and *Vibrio vulnificus* (SmcR) (4).

Because LuxR is responsible for reconfiguring gene expression as cells transition from LCD to HCD, it is a critical but complex global regulator. Previous studies used microarray, RNA-seq, and ChIP-seq analyses to show that LuxR binds to 115 promoters to control the expression of >400 genes (10, 11). Genes within the LuxR regulon are involved in a number of processes including bioluminescence, virulence, secretion, and metabolism (10, 12). The roles of some LuxR-regulated genes have been extensively characterized; however, many genes encode proteins that do not have predicted functions and are annotated as ‘hypothetical’ proteins. Here, we examined 16 LuxR-regulated genes that are predicted to be transcription factors. We determined the level of LuxR activation or repression of these genes and confirmed that LuxR directly binds to the promoters of 15 of these genes to directly control transcription regulation. Using RNA-seq, we describe the second tier of LuxR-regulated genes in *Vibrio harveyi*, which is comprised of 75 genes. These genes are associated with many different cellular processes discussed below.

## Results and Discussion

### LuxR regulates 16 transcription factors

Previous studies have determined the genes in the LuxR regulon in *V. harveyi* using microarray and RNA-seq analyses under near-identical conditions (10, 12). These analyses revealed 625 and 424 genes, respectively, that show significant ≥2-fold regulation (activation or repression) in the presence of LuxR. Among these LuxR-regulated genes are 16 that encode proteins with predicted transcription factor functionality. Based on the microarray and RNA-seq experiments, LuxR represses expression of 13 of these transcription factor genes and activates expression of 3 genes. To verify these results, we performed qRT-PCR to compare RNA levels of these transcription factors between wild-type and Δ*luxR* strains. As expected, the qRT-PCR results corroborate the microarray and RNA-seq expression data confirming that 13 genes are repressed while 3 genes are activated (Fig. 1). One gene, *00507*, was not significantly different between wild-type and Δ*luxR* in our qRT-PCR assay. However, in both the previous microarray and RNA-seq data, *00507* expression was found to be significantly different between the two strains (*p* < 0.0001) (11, 12). Therefore, we conclude that *00507* is activated by LuxR. Importantly, the fold-regulation by LuxR of each gene assayed by qRT-PCR closely mirrors the values determined using microarrays and RNA-seq (11, 12).

**Figure 1.**
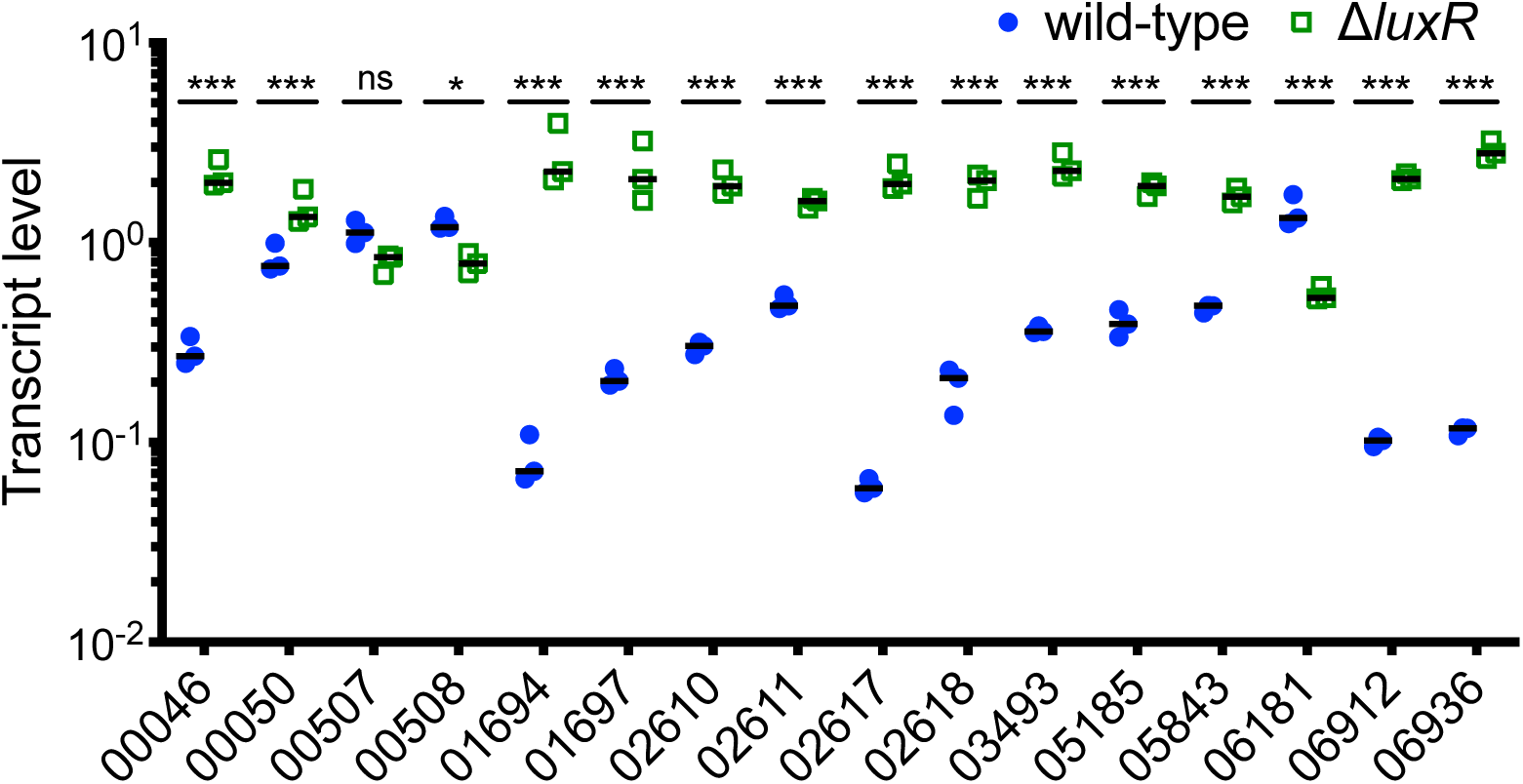
LuxR regulates 16 genes encoding putative transcription factors. Relative transcript levels of 16 genes determined by qRT-PCR from RNA isolated from wild-type (BB120) or Δ*luxR* (KM669) *V. harveyi* strains. Gene names are indicated under the graph and correspond to the GenBank annotation *VIBHAR_XXXXX*. Asterisks indicate significant differences between wild-type and Δ*luxR* strains (*, *p* <0.05; ***, *p* < 0.001; two-way analysis of variance (ANOVA) followed by Sidak’s multiple comparisons test on log-transformed data; *n*=3).

### LuxR directly binds to the promoter regions of 15 genes encoding transcription factors

Previous assays have shown that LuxR binds directly to several of the promoters for the 16 genes encoding transcription factors (10,11,13). LuxR ChIP-seq peaks were observed in the upstream regions of each of the 16 genes except *00508*, *02611*, *06912, 02618*, and the two type III secretion regulators, *exsA* and *exsB* (Fig. 2) (10, 14) However, electrophoretic mobility shift assays (EMSAs) showed that LuxR binds to the promoters of *exsA* and *exsB in vitro* (11, 15). It is not clear why LuxR ChIP data (from two studies) do not indicate LuxR binding at this locus but *in vitro* assays show LuxR is capable of binding (10, 14). One possibility is that LuxR does bind to this locus *in vivo* but at earlier time points in the growth curve. The two ChIP-seq datasets were performed at early and late stationary phase. Thus, it is possible that LuxR binds to the promoters of *exsA* and *exsB* in log-phase and is then outcompeted at stationary phase by another transcription factor. The lack of LuxR ChIP-seq peaks upstream of the *02611* and *00508* genes is not surprising because these genes are likely encoded in operons with *02610* and *00507*, respectively. The *02610-02611* and *00507-00508* gene pairs are located in close proximity in the same orientation, and RNA-seq data suggest that these genes are co-transcribed (12). Further, LuxR ChIP-seq peaks exist upstream of *02610* and *00507* (Fig. 2), indicating that LuxR directly regulates these two operons.

**Figure 2.**
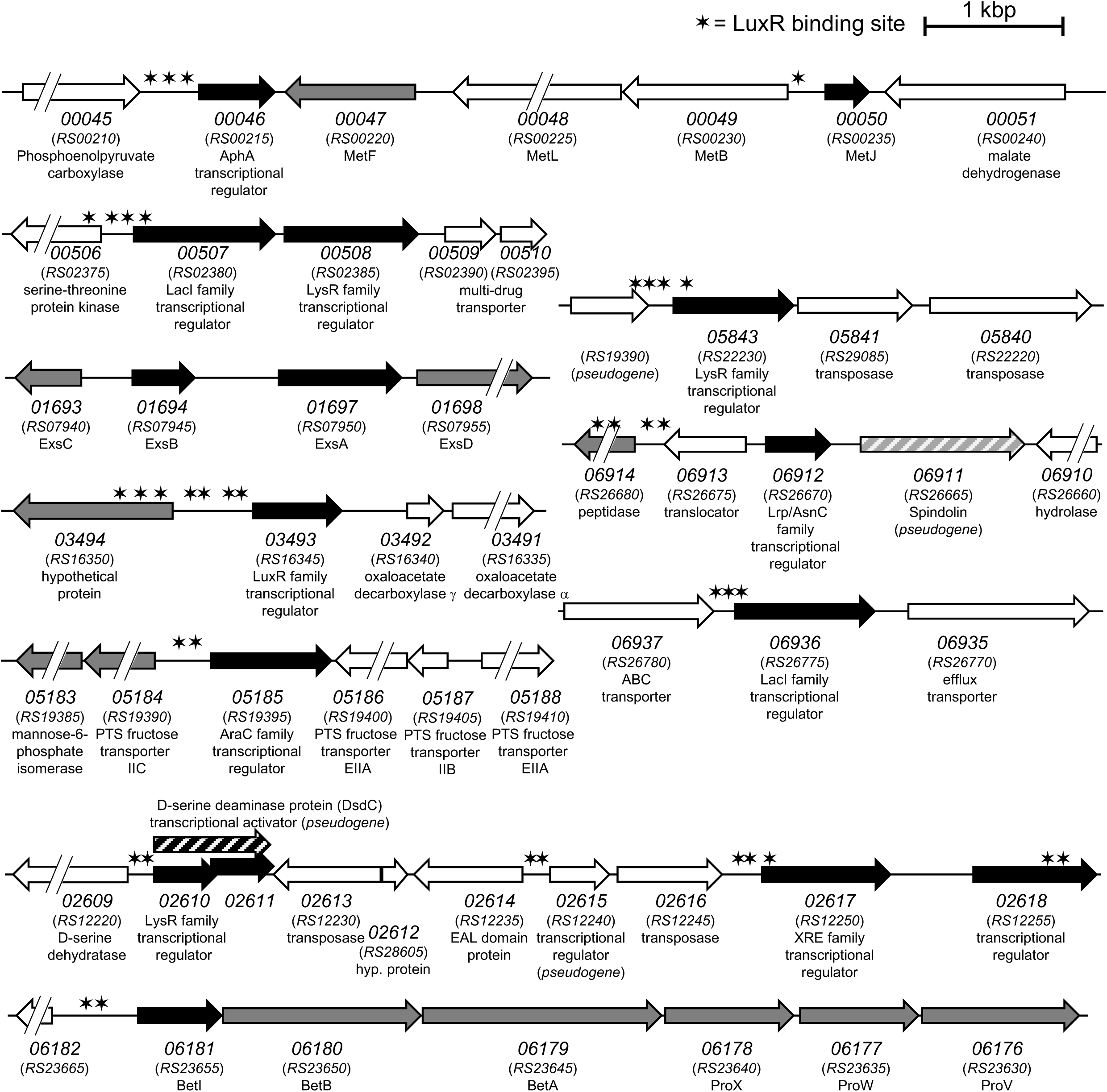
Genetic loci containing 16 putative transcription factor-encoding genes regulated by LuxR. Diagram showing the 16 loci containing genes encoding putative transcription factors (black arrows), and nearby genes regulated by LuxR >2-fold (gray arrows). Genes not regulated by LuxR >2-fold are shown as white arrows. Stars indicate locations of peaks in previously published ChIP-seq assays (10).

Using these previous data as a guide, we hypothesized that LuxR directly binds to the promoters of all of these 16 genes except *06912 and 02618*, the only genes for which no direct binding by LuxR has been previously observed. To investigate whether LuxR modulates the expression of these transcription factors directly or indirectly, we performed EMSAs with PCR-amplified DNA corresponding to the 400 bp upstream of the open reading frame (ORF) for each gene. LuxR binding was observed at all of the upstream loci tested except the region upstream of *06912* (Fig. 3). We did observe binding in the promoter of *02618 in vitro* and a minor peak in ChIP-seq, suggesting that LuxR has the capability of binding this promoter. Our results also confirmed that LuxR does not directly control *06912*. The dynamics of LuxR binding to these loci are variable. For example, some regions possess multiple LuxR binding sites (*e.g., 00507*) which is apparent from the super-shifted band. This is not uncommon, as there is an average of two LuxR binding sites per promoter in the LuxR regulon (10). Additionally, the relative affinity of LuxR binding sites within these promoters is variable. Some loci (*e.g., 06181*) are completely bound with as little as 10 nM LuxR while others required 100 nM LuxR protein to be shifted (*e.g., 01697*) (Fig. 3). Again, this is reminiscent of other LuxR-regulated operons (*e.g.*, *luxCDABE*) that show a range of LuxR binding affinities. Together, these results indicate that LuxR directly regulates 15 of the 16 genes encoding putative transcription factors.

**Figure 3.**
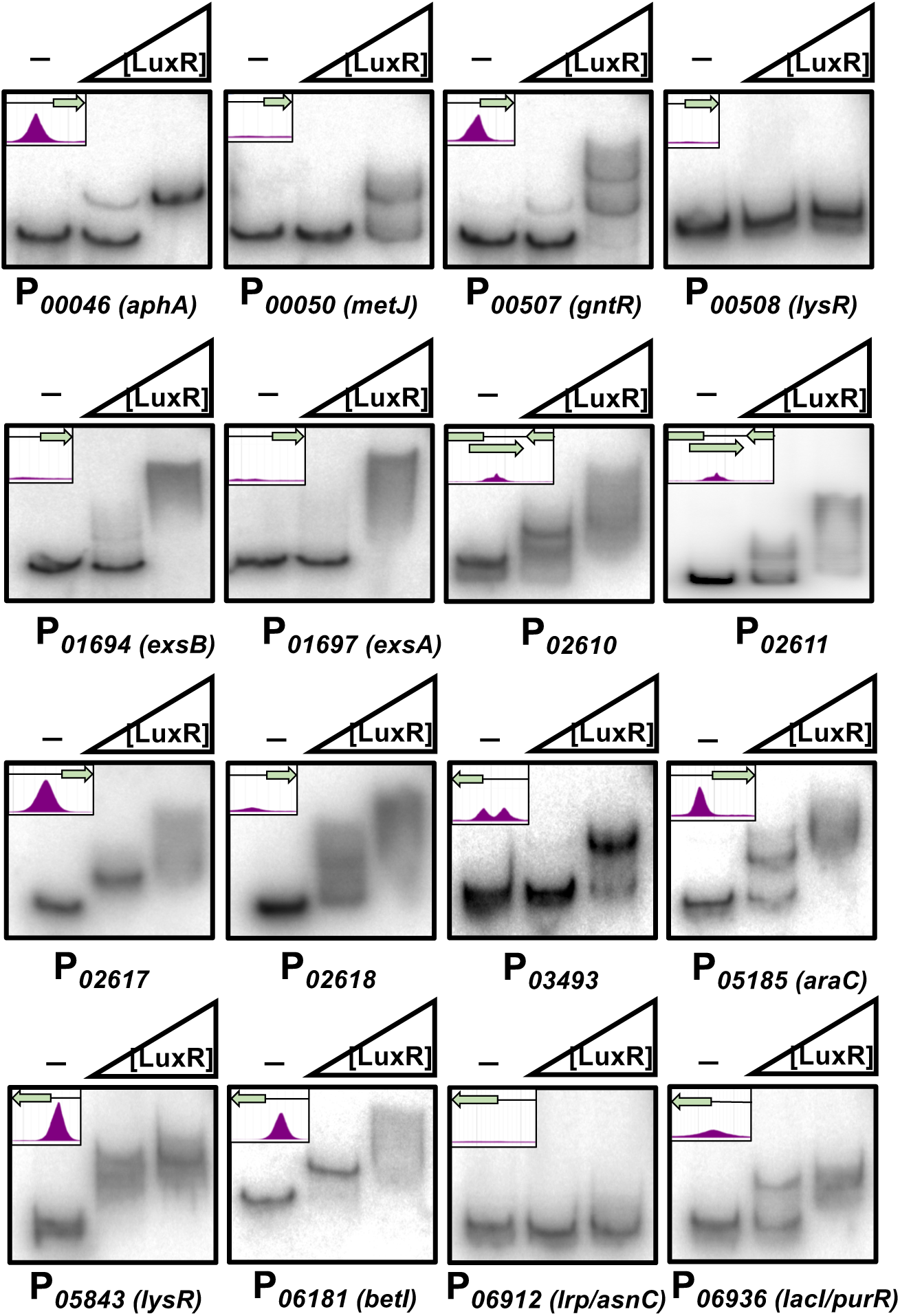
LuxR directly binds to the regions upstream of 14 genes encoding putative transcription factors. EMSAs containing PCR-generated, radiolabeled promoter DNA (400 bp) and purified LuxR protein at 0, 10, and 100 nM concentrations (except for P*_02617_* and P*_02618_* substrates in which 0, 20, and 100 nM were used). Insets show LuxR ChIP-seq peak profiles for each locus using data from a previous study (14).

### RNA-seq analysis determines the second tier of quorum-sensing regulation

Because ChIP-seq data indicate that LuxR directly regulates only ∼115 genes out of >400 genes in its regulon, we predicted that these 15 transcription factors control the remaining genes in the LuxR regulon. To examine this, we performed RNA-seq analyses on a subset of these genes to determine their regulons. We analyzed the regulon of the following genes predicted to encode transcription factors: *00050* (*metJ*), *02610-02611*, *02617*, *02618*, *05185*, *06912*, and *06936*. We did not investigate the other transcription factor genes as they have been previously characterized: 1) *06181* encodes BetI, which regulates the *betIBA-proXWV* operon in *V. harveyi* (6 genes) (13); 2) *01697* and *01694* encode ExsA and ExsB, respectively, which regulate the four type III secretion operons in *V. harveyi* (36 genes) (15); *00046* encodes AphA, which controls the LCD regulon in *V. harveyi* (167 genes) (3, 11). The remaining genes, *00507, 00508, 03493,* and *05843*, are regulated <3-fold by LuxR, thus we chose not to pursue them. Because each of the genes we chose to examine are repressed by LuxR, we used the Δ*luxR* background for all RNA-seq experiments to determine the regulon of each transcription factor.

#### VIBHAR_00050: MetJ

The *VIBHAR_00050* gene encodes the transcriptional regulator MetJ, which shares 80.7% amino acid identity with MetJ from *Escherichia coli*. Bacterial methionine biosynthesis has been extensively studied and the role of MetJ is well understood (16). In *E. coli*, MetJ function opposes a transcriptional activator, MetR, to repress many of the methionine biosynthetic enzymes including *metL*, *metA*, *metB*, *metC*, *metH*, and others (17–19). Although LuxR only represses the expression of *metJ* by 1.5-fold under the single condition we tested, we considered this regulation to be important because MetR regulates one of the most highly activated quorum-sensing phenotypes, bioluminescence, in *V. harveyi* (20).

Among all of the transcription factors we investigated, deletion of *metJ* affected the largest number of genes; our RNA-seq experiment revealed that *metJ* controls the expression of 49 genes (+/− 2-fold, *p*-adjusted ≤ 0.05, Table 1). MetJ activates the expression of 9 genes while repressing the remaining 40. The degree of regulation exhibited by MetJ is heavily shifted towards repression as the most highly activated gene in its regulon is only upregulated by ∼4-fold while the *met* genes (the most highly repressed genes) are downregulated ∼15- to 75-fold. These findings are in strong agreement with published literature that describes MetJ as a repressor of the methionine biosynthetic genes (17, 18). In addition to the methionine biosynthesis genes, MetJ regulates the expression of several other metabolic enzymes, suggesting that this transcription factor fine-tunes other sectors of cellular metabolism. At HCD, *V. harveyi* maximally produces LuxR, repressing the expression of MetJ. Thus, the methionine biosynthetic genes are upregulated at HCD. One possible explanation for this finding is linked to the fact that AI-2 production is dependent on methionine and cysteine production (21). As the population reaches stationary phase and nutrient availability decreases, it may be important for cells to increase expression of the methionine biosynthesis proteins to maintain AI-2 production. This compensatory mechanism would ensure that AI-2 mediated communication between cells is uninterrupted as local nutrient availability changes.

**Table 1.** Genes regulated by VIBHAR_00050 (MetJ).

| Locus ID | Old locus ID | Fold regulation | p-adjusted value | Predicted function |
| --- | --- | --- | --- | --- |
| VIBHAR_RS25170 | VIBHAR_06544 | 4.0 | 7.5E-07 | hypothetical protein |
| VIBHAR_RS23080 | VIBHAR_06041 | 3.8 | 4.6E-02 | hypothetical protein |
| pspA | VIBHAR_01872 | 3.1 | 1.5E-33 | phage shock protein PspA |
| pspB | VIBHAR_01873 | 2.9 | 3.8E-21 | envelope stress response membrane protein PspB |
| pspC | VIBHAR_01874 | 2.4 | 3.2E-28 | PspC domain-containing protein |
| VIBHAR_RS00035 | VIBHAR_00008 | 2.4 | 9.3E-04 | envelope stress response protein PspG |
| VIBHAR_RS12360 | VIBHAR_02639 | 2.1 | 6.4E-10 | TetR/AcrR family transcriptional regulator |
| VIBHAR_RS13045 | VIBHAR_02785 | 2.0 | 1.3E-07 | hypothetical protein |
| VIBHAR_RS18605 | VIBHAR_05000 | 2.0 | 4.0E-04 | glycine C-acetyltransferase |
| VIBHAR_RS25745 | VIBHAR_06685 | -2.0 | 2.8E-03 | hypothetical protein |
| VIBHAR_RS20725 | VIBHAR_05495 | -2.1 | 3.1E-05 | protoheme IX farnesyltransferase 1 |
| VIBHAR_RS07970 | VIBHAR_01701 | -2.1 | 2.6E-10 | EscD/YscD/HrpQ family type III secretion system inner membrane ring protein |
| VIBHAR_RS07990 | VIBHAR_01705 | -2.1 | 3.8E-10 | type III export protein |
| VIBHAR_RS06990 | VIBHAR_01482 | -2.1 | 3.7E-02 | hypothetical protein |
| VIBHAR_RS10100 | VIBHAR_02153 | -2.1 | 1.9E-04 | hypothetical protein |
| VIBHAR_RS23730 | VIBHAR_06196 | -2.2 | 8.2E-03 | pilus assembly protein CpaC |
| VIBHAR_RS10700 | VIBHAR_02279 | -2.2 | 1.0E-03 | formate dehydrogenase |
| VIBHAR_RS19920 | VIBHAR_05304 | -2.2 | 1.5E-02 | 1,4-alpha-glucan-branching protein |
| VIBHAR_RS21930 | VIBHAR_05775 | -2.2 | 2.4E-02 | hypothetical protein |
| VIBHAR_RS23735 | VIBHAR_06197 | -2.3 | 4.0E-02 | pilus assembly protein CpaB |
| cyoC | VIBHAR_05493 | -2.3 | 1.2E-07 | cytochrome o ubiquinol oxidase subunit III |
| VIBHAR_RS21180 | VIBHAR_05607 | -2.4 | 1.3E-03 | hypothetical protein |
| VIBHAR_RS10105 | VIBHAR_02154 | -2.4 | 1.0E-05 | hypothetical protein |
| VIBHAR_RS01205 | VIBHAR_00256 | -2.4 | 2.3E-10 | glutamate racemase |
| VIBHAR_RS23745 | VIBHAR_06199 | -2.5 | 5.2E-03 | Flp family type IVb pilin |
| VIBHAR_RS22625 | VIBHAR_05936 | -2.5 | 4.2E-02 | methyl-accepting chemotaxis protein |
| VIBHAR_RS10705 | VIBHAR_02280 | -2.5 | 1.1E-05 | formate dehydrogenase subunit gamma |
| VIBHAR_RS16695 | VIBHAR_RS16695 | -2.7 | 4.6E-05 | hypothetical protein |
| VIBHAR_RS18715 | VIBHAR_05026 | -3.3 | 4.6E-05 | phage capsid protein |
| VIBHAR_RS23740 | VIBHAR_06198 | -3.5 | 7.1E-03 | prepilin peptidase CpaA/TadV |
| VIBHAR_RS12030 | VIBHAR_02564 | -3.5 | 6.2E-06 | sodium transporter |
| VIBHAR_RS01210 | VIBHAR_00257 | -4.9 | 1.9E-14 | hypothetical protein |
| VIBHAR_RS17385 | VIBHAR_03712 | -6.2 | 6.4E-238 | methionine synthase |
| VIBHAR_RS01215 | VIBHAR_00260 | -6.3 | 7.1E-06 | hypothetical protein |
| VIBHAR_RS22555 | VIBHAR_05920 | -6.9 | 2.3E-04 | DUF1097 domain-containing protein |
| VIBHAR_RS05235 | VIBHAR_01104 | -8.4 | 3.3E-23 | sodium:proton antiporter |
| VIBHAR_RS01220 | VIBHAR_00261 | -10.2 | 5.8E-195 | vitamin B12 transporter BtuB |
| VIBHAR_RS12990 | VIBHAR_02773 | -15.6 | 3.1E-23 | HTH-type transcriptional regulator MetR |
| metK | VIBHAR_03568 | -17.1 | 0.0E+00 | S-adenosylmethionine synthase |
| metN | VIBHAR_01202 | -19.0 | 2.3E-233 | D-methionine ABC transporter, ATP-binding protein |
| VIBHAR_RS11700 | VIBHAR_02491 | -26.8 | 0.0E+00 | homoserine O-succinyltransferase |
| metQ | VIBHAR_01200 | -28.9 | 0.0E+00 | methionine ABC transporter substrate-binding protein MetQ |
| VIBHAR_RS05680 | VIBHAR_01201 | -31.1 | 5.2E-151 | ABC transporter permease |
| VIBHAR_RS13115 | VIBHAR_02800 | -44.2 | 2.2E-20 | 5-methyltetrahydropteroyltriglutamate-homocysteine methyltransferase |
| metF | VIBHAR_00047 | -45.2 | 0.0E+00 | methylenetetrahydrofolate reductase |
| metL | VIBHAR_00048 | -49.6 | 0.0E+00 | bifunctional aspartate kinase/homoserine dehydrogenase II |
| metB | VIBHAR_00049 | -56.2 | 0.0E+00 | cystathionine gamma-synthase |
| VIBHAR_RS13120 | VIBHAR_02801 | -64.7 | 2.8E-30 | DUF1852 domain-containing protein |
| VIBHAR_RS12985 | VIBHAR_02772 | -76.8 | 1.2E-122 | 5-methyltetrahydropteroyltriglutamate-homocysteine methyltransferase (MetE) |

Distinct from MetJ, MetR is a direct repressor of bioluminescence in *V. harveyi* (22). Consistent with data from *E. coli*, MetJ in *V. harveyi* represses the expression of MetR (23). To investigate the effects of this regulatory loop, we assayed light production in the Δ*metJ* strain (Fig. 4A). The wild-type strain BB120 was used as a reference and shows increased bioluminescence as a function of cell growth. Importantly, light production requires LuxR, and we note that the Δ*luxR* strains do not bioluminesce. Compared to the wild-type strain, the Δ*metJ* strain exhibits a ∼2-fold reduction in bioluminescence at HCD. To verify that MetJ is responsible for this difference, we attempted to complement the Δ*metJ* strain using an IPTG-inducible expression vector. To our surprise, when MetJ is expressed in *trans* in the Δ*metJ* strain, bioluminescence is not restored to wild-type levels (Fig. S1). Typically, this indicates the presence of a polar mutation in the parent strain that is affecting bioluminescence. However, we observed complementation of another MetJ-regulated gene, *metL* (*VIBHAR_00048*), via qRT-PCR (Fig. 4B). These results suggest that the Δ*metJ* strain does not harbor a polar mutation that affects all the genes in the MetJ regulon. It is thus far not clear how the Δ*metJ* mutation is affecting bioluminescence expression, but the result is consistent through multiple biological experiments. The *metJ* gene is not in an operon and rather is flanked by genes in opposing directions (Fig. 2), but perhaps there is an effect on the transcriptional initiation and/or termination of the adjacent gene(s) in the mutant strain.

**Figure 4.**
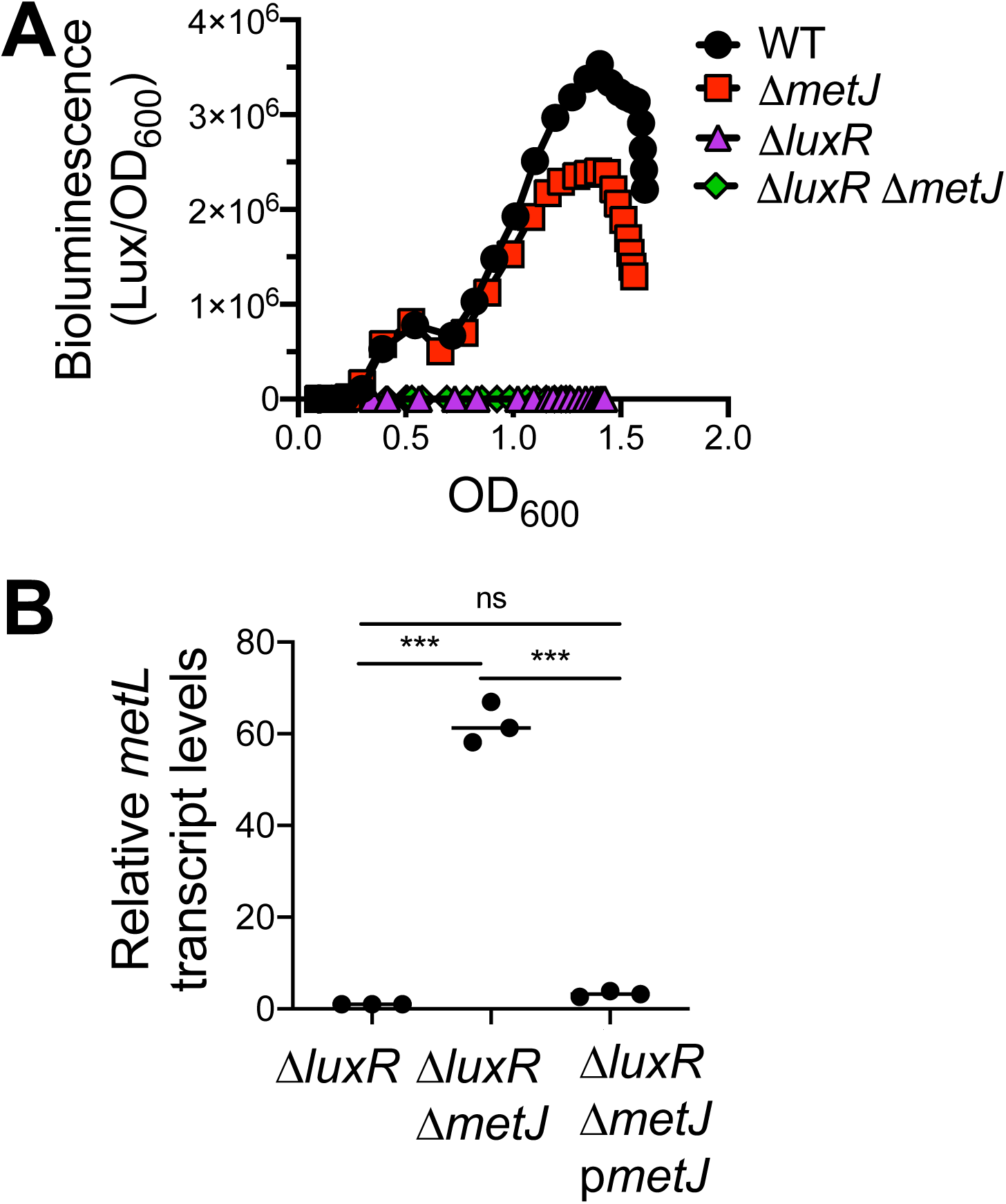
MetJ regulates bioluminescence and methionine synthesis. (A) Bioluminescence (lux/OD_600_) produced by wild-type (BB120), Δ*luxR* (KM669), Δ*metJ* (RRC176), and Δ*luxR* Δ*metJ* (RRC178) strains. Data shown are representative of four independent experiments. (B) Transcript levels of *metL* (*VIBHAR_00048*) determined by qRT-PCR of RNA isolated from Δ*luxR* + empty vector (AB087), Δ*metJ* + empty vector (AB125), and Δ*metJ* + p*metJ* (AB121) strains. 50 µM IPTG was added to all three strains. One way analysis of variance (ANOVA) followed by Tukey’s multiple comparisons test was performed (*n* = 3, *p* < 0.001).

Although MetJ activates the expression of only 9 genes, some of these are predicted to be involved in interesting behaviors. For example, *VIBHAR_02639* encodes a TetR/AcrR family transcriptional regulator, which typically represses genes responsible for modifying, degrading, and/or exporting antibiotics (24, 25). Binding of the drug by a TetR/AcrR-type protein relieves repression of antibiotic tolerance genes, and the cytotoxic effects of the drug are mitigated (26). Structural homology analysis performed with Phyre2 identified the closest homologue of VIBHAR_02639 as PsrA from *Pseudomonas aeruginosa* (39% amino acid identity, 98% coverage). PsrA, another TetR/AcrR-type protein, controls the expression of type III secretion, RpoS, swarming motility, and biofilm formation in *P. aeruginosa* (27–29). PsrA responds to long-chain fatty acids as a means to control expression of fatty acid degradation genes (*fad* operon) (30, 31). In *V. harveyi*, this PsrA-like gene is located directly upstream of a gene annotated as an acyl-CoA dehydrogenase (*VIBHAR_02638*). Acyl-CoA dehydrogenase enzymes, such as FadE from *E. coli*, perform the initial steps of the β-oxidation cycle of fatty acid metabolism. Indeed, VIBHAR_02638 shares significant homology to FadE (50% amino acid identity, 90% coverage). Thus, based on genetic organization and protein homology, the VIBHAR_02638/39 operon that is activated by MetJ may be involved in fatty acid metabolism. The net effect of LuxR on this operon is negative; as cells reach HCD LuxR represses MetJ, which activates VIBHAR_02628/39, ultimately yielding a decrease in the expression of these genes at HCD (observed in (12)). The most likely rationale for this regulatory mechanism is that stationary populations have lower energy demands compared to actively growing cells. Thus, the QS of *V. harveyi* likely plays a role in fine-tuning of lipid metabolism as nutrient availability changes.

#### VIBHAR_02610-2611: DsdC

Genes *02610* and *02611* were first annotated by Genbank as two separate ORFs, with *02611* using a TTG start codon and both having homology to the LysR-type D-serine deaminase transcriptional activator DsdC (Fig. 2). In agreement with this predicted function is the observation that directly upstream is a gene that encodes a homologue of DsdA (D-serine dehydratase); this mirrors the genetic organization of *dsdC/A* in various *E. coli* strains (32). However, the most recent Genbank annotation of the *V. harveyi* BB120 genome predicts that this region encodes a pseudogene because it does not recognize the TTG start and instead denotes a frameshift. Because GeneMarkS (33) also predicts two open reading frames and other genes encoded by *V. harveyi* use a TTG start codon, we analyzed this region as containing two ORFS (*02610* and *02611*) and constructed deletion mutants of each gene individually.

Our RNA-seq experiments revealed that VIBHAR_02610 controls the expression of 13 genes (+/− 2-fold, *p*-adjusted ≤ 0.05, Table 2). Among these 13 genes, 4 are repressed by VIBHAR_02610 while 9 are activated. The 4 genes repressed by VIBHAR_02610 appear to be involved in acetate transport/processing, conjugal DNA transfer, and metal transport. VIBHAR_02610 represses the expression of an acetate symporter (VIBHAR_00160) and an acetate-CoA ligase (VIBHAR_00169). Presumably, these proteins import acetate and convert it to acetate-CoA (actyl-CoA), which can be fed directly into the TCA cycle. Because these genes are repressed by VIBHAR_02610, which is repressed by LuxR, the expression of these genes is likely elevated at HCD. This hypothesis holds true for at least *VIBHAR_00169* whose expression is 1.7-fold higher in the Δ*luxR* strain (12).The increased expression of genes involved in acetate metabolism aligns well with previous findings that show QS is utilized by *V. cholerae* to detoxify organic acids that accumulate in matured populations (34).

**Table 2.** Genes regulated by VIBHAR_02610

| Locus ID | Old locus ID | Fold regulation | p-adjusted value | Predicted function |
| --- | --- | --- | --- | --- |
| VIBHAR_RS25170 | VIBHAR_06544 | 3.0 | 1.6E-04 | hypothetical protein |
| VIBHAR_RS12355 | VIBHAR_02638 | 2.9 | 3.5E-08 | acyl-CoA dehydrogenase |
| VIBHAR_RS00035 | VIBHAR_00008 | 2.7 | 3.0E-04 | envelope stress response protein PspG |
| VIBHAR_RS12360 | VIBHAR_02639 | 2.5 | 8.9E-15 | TetR/AcrR family transcriptional regulator |
| pspA | VIBHAR_01872 | 2.3 | 1.6E-16 | phage shock protein PspA |
| VIBHAR_RS03960 | VIBHAR_00836 | 2.2 | 7.0E-08 | hypothetical protein |
| pspB | VIBHAR_01873 | 2.2 | 1.1E-10 | envelope stress response membrane protein PspB |
| VIBHAR_RS13960 | VIBHAR_02991 | 2.2 | 4.7E-02 | ribonuclease T |
| pspC | VIBHAR_01874 | 2.1 | 1.7E-18 | PspC domain-containing protein |
| acs | VIBHAR_00169 | -3.2 | 1.7E-03 | acetate--CoA ligase |
| VIBHAR_RS00780 | VIBHAR_00160 | -3.6 | 4.0E-02 | cation acetate symporter |
| VIBHAR_RS27820 | VIBHAR_p08218 | -7.2 | 1.1E-03 | TraY domain-containing protein |
| VIBHAR_RS00775 | VIBHAR_00159 | -9.7 | 3.0E-02 | DUF4212 domain-containing protein |

Regarding conjugal DNA transfer, VIBHAR_02610 represses *VIBHAR_p08218*, which is annotated as a TraY-domain containing protein. In support of this annotation, VIBHAR_p02818 shares an appreciable amount of homology (43% amino acid identity, 53% coverage) with TraY from *E. coli*. Adding to this, the *VIBHAR_p08218* gene is located on the naturally occurring plasmid in *V. harveyi* and is organized in an operon with other genes carrying the *tra* annotation. The Tra proteins have been extensively studied and it is now clear that TraY plays a critical role in assembling the relaxasome, a protein complex that facilitates plasmid transfer, at the origin of transfer (35). As LuxR is a repressor of VIBHAR_02610, VIBHAR_p08218 is activated at HCD. This agrees with current models in which bacterial conjugal DNA transfer is stimulated by high population densities and QS (36, 37).

The gene most highly repressed by VIBHAR_02610 is *VIBHAR_00159*, which is annotated as encoding a hypothetical protein containing a DUF4212 domain. Using BLAST and Phyre2 to probe the potential function of this protein, we discovered secondary structure prediction homology to *E. coli* protein ZntB. In *E. coli*, this membrane-embedded protein oligomerizes to form a membrane channel that is used to transport zinc into the cytoplasm (38). The homology between ZntB and VIBHAR_00159 lies within the two transmembrane helices – this is meaningful because these helices are directly involved in the transport of zinc through the membrane. Interestingly, ZntB is a 327-amino acid protein while VIBHAR_00159 is only 87 amino acids in length; this observation suggests that VIBHAR_00159 is a truncated version of ZntB. It is possible that, in *V. harveyi*, VIBHAR_00159 alone is not sufficient to transport zinc but by associating with an additional protein(s), can form a complex that is zinc transport competent. Zinc uptake is critically important for pathogenic bacteria as many metalloproteases require zinc to function (39). Our data implies that at least one possible zinc transport protein is upregulated as cells reach HCD, which is when virulence gene expression programs are activated in other *vibrios* such as *V. parahaemolyticus* and *V. vulnificus* (40, 41).

Genes activated by VIBHAR_02610 are associated with membrane stress response, transcription, and RNA processing. The membrane stress response genes correspond to the <u>p</u>hage <u>s</u>hock <u>p</u>roteins *pspABCG* – these proteins respond to cell envelope stresses through a variety of mechanisms (42, 43). Interestingly, the *psp* genes and the PsrA-like transcription factor (VIBHAR_02639) are all activated by both VIBHAR_02610 and MetJ. It is possible that VIBHAR_02610 and MetJ function together to coordinate certain gene expression programs.

Another interesting gene that is activated by VIBHAR_02610 is RNase T. RNase T is a 3-5’ exoribonuclease that is critically important for the maturation of stable RNAs (44). It is essential for the maturation of 5S and 23S rRNA as well as tRNA turnover (45). Our data show that VIBHAR_02610 activates the expression of RNase T ∼2.2-fold. Not surprisingly, VIBHAR_02610 is also responsible for activating the expression of many 16S/23S rRNAs and multiple tRNAs, however it is important to note that these genes are only present in the regulon when the *p*-adjusted value cutoff is raised to 0.07-0.12. Because LuxR represses VIBHAR_02610, it is probable that the production of these rRNAs/tRNAs is diminished at HCD. This hypothesis fits well with the general physiology; as cells transition from exponential growth phase into stationary phase (when LuxR is maximally expressed), transcription and translation rates are reduced (46). Thus, *V. harveyi* may reduce global translation via LuxR-mediated repression of VIBHAR_02610. A protein expression profile of VIBHAR_02610 and correlative translation rate (or growth rate) experiments would provide more insight on this hypothesis.

In stark contrast to VIBHAR_02610, VIBHAR_02611 regulates the expression of only 2 genes in *V. harveyi* (+/− 2-fold, *p*-adjusted ≤ 0.05, Table 3). Both of these genes are repressed ∼3-fold by VIBHAR_02611. VIBHAR_01640 is annotated as a hypothetical protein; however, BLAST analysis indicates that the protein product of this gene possesses a helix-turn-helix domain which is common among transcriptional regulators (47). This could explain why VIBHAR_02611 regulates so few genes as it is part of a transcriptional regulatory cascade. Additionally, this gene is organized at the head of a 3-gene operon. Within this operon is a gene that encodes a hypothetical protein that possesses homology to an RNA-directed DNA polymerase, or reverse transcriptase, in *Salmonella enterica* (32% amino acid identity, 93% coverage). Unfortunately, the biological function of bacterial reverse transcriptases has remained enigmatic since their discovery in 1970 (48). From the small amount of literature that exists on this topic, it is apparent that various genera of bacteria (*Escherichia*, *Myxococcus*, and *Stigmatella*) encode a chromosomal copy of a reverse transcriptase that is involved in the production of multi-copy singled-stranded DNA, or msDNA (49, 50). Little is known regarding the function of msDNA however, it has been reported that msDNA can be transmitted between cells in a population (51). Interestingly, the formation of mature msDNA involves the activity of RNaseH. The other gene that is regulated by VIBHAR_02611 is *VIBHAR_06361*, which encodes an endo/exonuclease and could fulfill the role of RNaseH. Furthermore, within this chromosomal locus, we observe LuxR-dependent transcripts that are produced from the non-coding strand; this could represent the RNA template that is utilized by the reverse transcriptase to produce msDNA. Further dissection of this locus to explore its functionality will undoubtedly yield informative results.

**Table 3.** Genes regulated by VIBHAR_02611

| Locus ID | Old locus ID | Fold regulation | p-adjusted value | Predicted function |
| --- | --- | --- | --- | --- |
| VIBHAR_RS07715 | VIBHAR_01640 | -2.8 | 1.41E-02 | hypothetical protein |
| VIBHAR_RS24415 | VIBHAR_06361 | -2.9 | 9.10E-03 | endonuclease/exonuclease/phosphatase family protein |

#### *VIBHAR_02617*: XRE family transcription factor and *VIBHAR_02618*

VIBHAR_02617 is classified as an XRE-type transcriptional regulator. These transcriptional regulators are proteins that contain a helix-turn-helix DNA binding motif and are homologous to the CI and Cro proteins from bacteriophage λ. VIBHAR_02618 does not have an obvious classification and instead is annotated generally as a transcriptional regulator. We performed RNA-seq using the respective Δ*VIBHAR_02617* and Δ*VIBHAR_02618* strains and observed unexpected results. First, the regulons for each of these regulators are very small; VIBHAR_02617 regulates 3 genes while VIBHAR_02618 regulates 2 genes (+/− 2-fold, *p*-adjusted ≤ 0.05, Table 4 and Table 5). Second, both genes regulated by VIBHAR_02618 are also regulated by VIBHAR_02617, and in the same manner (both upregulated). Interestingly, the two co-regulated genes (*VIBHAR_07127* and *VIBHAR_05996*) encode proteins that are >94% identical in amino acid sequence. Unfortunately, these genes encode proteins with no clear homologues discernable via BLAST/Phyre2; their functions will need to be determined experimentally. We investigated the putative promoter regions upstream of both of these genes and found them to be nearly identical (99% conservation up to −300 bp from the start codon). These observations suggest that these ORFs likely originated from a duplication event. The exquisite conservation of not only amino acid sequence but promoter sequence as well prompts the hypothesis that these proteins serve a critical function.

**Table 4.** Genes regulated by VIBHAR_02617

| Locus ID | Old locus ID | Fold regulation | p-adjusted value | Predicted function |
| --- | --- | --- | --- | --- |
| VIBHAR_RS27540 | VIBHAR_07127 | 2.4 | 4.6E-15 | hypothetical protein |
| VIBHAR_RS22885 | VIBHAR_05996 | 2.3 | 3.4E-07 | hypothetical protein |
| VIBHAR_RS01705 | VIBHAR_00366 | 2.1 | 3.9E-03 | phosphomethylpyrimidine synthase (ThiC) |

**Table 5.** Genes regulated by VIBHAR_02618

| Locus ID | Old locus ID | Fold regulation | p-adjusted value | Predicted function |
| --- | --- | --- | --- | --- |
| VIBHAR_RS27540 | VIBHAR_07127 | 3.5 | 1.2E-27 | hypothetical protein |
| VIBHAR_RS22885 | VIBHAR_05996 | 3.3 | 1.3E-13 | hypothetical protein |

The gene that is regulated by VIBHAR_02617 but not VIBHAR_02618 is *VIBHAR_00366*, which is annotated as a phosphomethylpyrimidine synthase. Several BLAST searches revealed this protein as ThiC, an important member of the thiamine biosynthetic pathway in bacteria (52–54). Details regarding thiamine biosynthesis are explored in the proceeding section. VIBHAR_02617 activates the expression of ThiC; thus, LuxR effectively represses this arm of thiamine biosynthesis.

#### *VIBHAR_05185*: AraC family transcription factor

The protein encoded by *VIBHAR_05185* is annotated as an AraC-type transcriptional regulator. AraC in *E. coli*, and homologues from other organisms, activates the expression of the arabinose utilization operons, *araBAD/FGH/E*, upon binding arabinose (55). Our RNA-seq data indicates that VIBHAR_05185 regulates 5 genes in *V. harveyi* (+/− 2-fold, *p*-adjusted ≤ 0.05, Table 6) that are involved in thiamine biosynthesis, fructose transport, and c-di-GMP degradation. The genes involved in thiamine biosynthesis, namely *thiH*, *thiE*, and *thiF*, are all repressed by VIBHAR_05185. In fact, the entire *thiCEFSGH* operon is repressed by VIBHAR_05185 but *thiC*, *thiS*, and *thiG* fall slightly below our 2-fold cutoff. The literature regarding the regulation of the *thi* genes is largely focused on the posttranscriptional level in which the THI-box riboswitch serves as a negative feedback control point (56). A transcriptional regulator that modulates the expression of the *thi* genes has yet to be identified in bacteria. A regulator has been identified in Archaea but it is not homologous to VIBHAR_05185 (57). It is possible that VIBHAR_05185 directly represses expression of the *thi* operon in *V. harveyi*. Conversely, and equally as likely, it is possible that secondary effects produced by VIBHAR_05185 (i.e. regulation of other genes/cellular processes) influence expression of this operon. Regardless, the net effect is that thiamine biosynthesis is upregulated as the population reach quorum. Thiamine and its derivatives function as co-factors for many cellular enzymes (58, 59). Thus, it is probable that QS fine-tunes the production of thiamine to accommodate the enzymatic activity of a growing population.

**Table 6.**
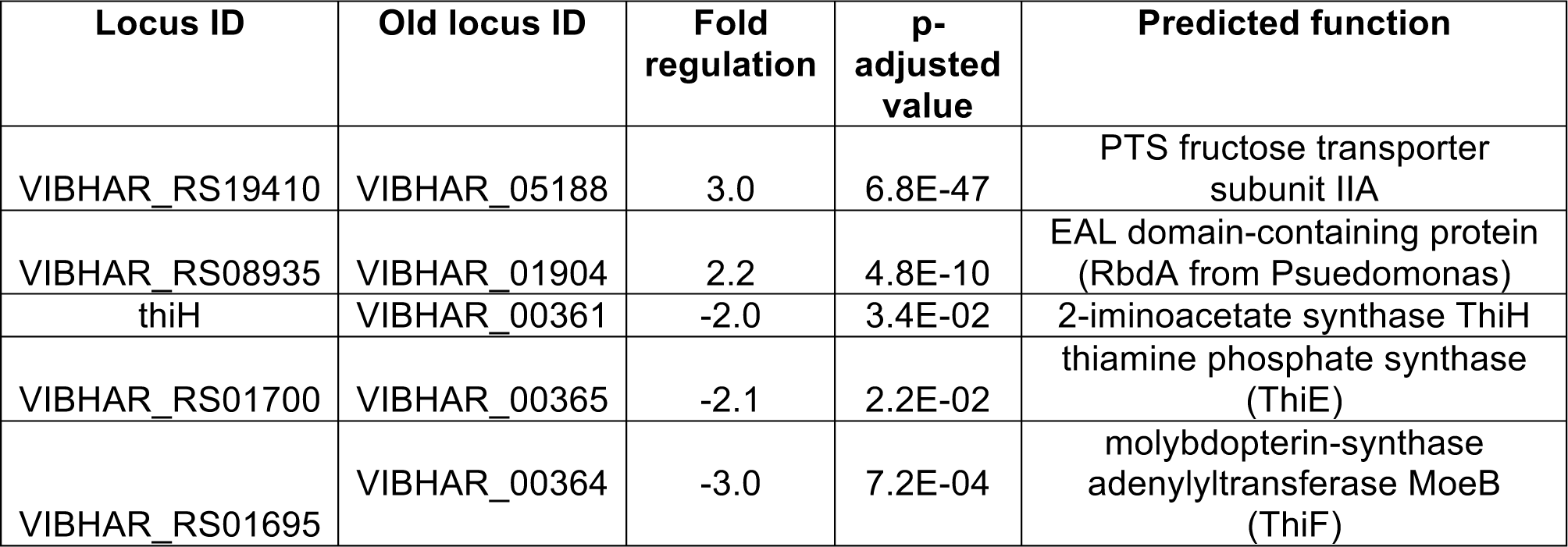
Genes regulated by VIBHAR_05185

In addition to the *thi* operon, VIBHAR_05185 activates the expression of a fructose transporter, encoded just downstream of *VIBHAR_05185* (Fig. 2). Because the expression of the fructose transporter gene is activated by VIBHAR_05185, which is repressed by LuxR, QS effectively reduces the expression of this gene at HCD. Consistent with this hypothesis, LuxR represses other components of the fructose PTS system that neighbor *VIBHAR_05185* at this locus (12). Interestingly, the media used in this experiment (LB with added salt) contains < 100 μM sugars (60). In other media with fructose, gene expression changes might be more substantial throughout quorum-sensing stages. We surmise that as populations of *V. harveyi* mature and reach HCD states, the availability of fructose likely decreases substantially. Thus, QS serves to shut down elements of fructose uptake as the carbon source is depleted.

Another interesting constituent of the VIBHAR_05185 regulon is VIBHAR_01904. This gene encodes an EAL-domain containing protein and further analysis using Phyre2 predicts significant secondary structure homology to RbdA from *P. aeruginosa*. RbdA is a PAS-GGDEF-EAL protein that negatively regulates biofilm formation while simultaneously stimulating dispersal (61, 62). Although RbdA contains both diguanylate cyclase and phosphodiesterase motifs, it is primarily responsible for degrading c-di-GMP *in vivo*. In addition to modulating intracellular concentrations of c-di-GMP, the PAS domain of RbdA serves as an O_2_ sensor (61). It has been postulated that RbdA is a component of low-oxygen stress response in *Pseudomonas*. When comparing the homology between RbdA and VIBHAR_01904, residues are mostly conserved in the PAS and EAL domains. The EAL domain is more strongly conserved so it is likely that VIBHAR_01904 also functions as a phosphodiesterase *in vivo*. Conversely, the PAS domain shows weaker conservation – this could be explained by a difference in ligand identity. Careful and thorough investigation of this protein could provide insight on c-di-GMP regulation and its connection to QS in *V. harveyi*.

#### *VIBHAR_06912*: Lrp/AsnC family transcription factor

The protein encoded by *VIBHAR_06912* belongs to the Lrp/AsnC family of transcription factors. These proteins are equipped with a DNA binding domain to interact with promoter DNA as well as a ligand binding domain to sense a small molecule/metabolite (*e*.*g*., leucine in the case of Lrp) (63). We were surprised to find that VIBHAR_06912 has no significant effect on gene expression in *V. harveyi* (+/− 2-fold, *p*-adjusted value ≤ 0.05). We rationalized that, because this transcription factor probably binds a ligand, it is possible that the regulon is only controlled under certain environmental conditions that differ from our laboratory conditions. This transcription factor is well conserved in other vibrios including *V. parahaemolyticus* as well as other bacteria such as *Ferrimonas*, *Shewanella*, and *Bacterioplanes* (>77% amino acid identity, 100% coverage). These organisms share one distinct property – they are all marine bacteria. This observation supports the idea that VIBHAR_06912 may be utilized under specific conditions and that these conditions are experienced by *Vibrio*, *Ferrimonas*, *Shewanella*, and *Bacterioplanes* alike.

#### *VIBHAR_06936*: LacI family transcription factor

The protein encoded by *VIBHAR_06936* belongs to the LacI/MalR transcription factor family. In *V. harveyi*, this transcription factor regulates 1 gene (+/− 2-fold, p-adjusted ≤ 0.05, Table 7). Interestingly, this gene is *thiC*, which is also regulated by VIBHAR_05185. Similar to VIBHAR_05185, *thiE* and *thiF* are also regulated by VIBHAR_06936, however they fall slightly below our 2-fold change in gene expression cutoff. Furthermore, VIBHAR_06936 and VIBHAR_05185 have opposite regulatory effects on *thiCEF*; VIBHAR_06936 activates expression while VIBHAR_05185 represses it. The degree of regulation of *thiCEF* by VIBHAR_05185/06936 is very similar – both proteins exert ∼2-fold change in expression. This means that the net change in expression is negated by the opposing action of these regulators. However, *thiSGH*, which are regulated by VIBHAR_05185 but not VIBHAR_06936, would remain differentially regulated in wild-type cells. Thus, it is conceivable that by modulating specific parts of the thiamine biosynthetic pathway, QS in *V. harveyi* is capable of precisely controlling metabolic flux through this pathway; this may be important for redirecting precursor molecules into alternative pathways.

**Table 7.**
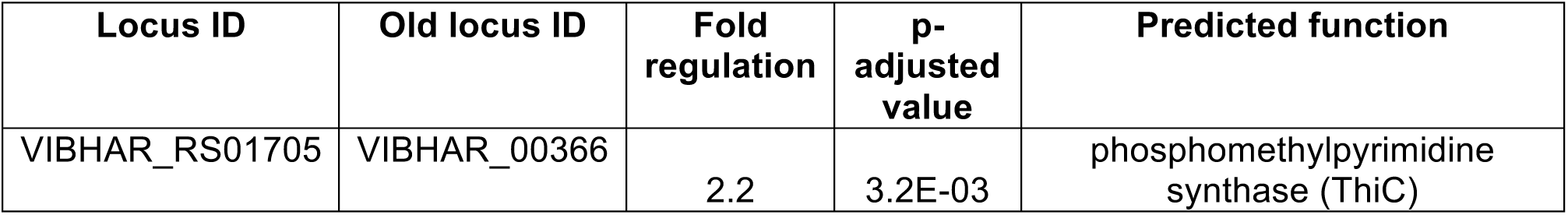
Genes regulated by VIBHAR_06936

| Locus ID | Old locus ID | Fold regulation | p-adjusted value | Predicted function |
| --- | --- | --- | --- | --- |
| VIBHAR_RS01705 | VIBHAR_00366 | 2.2 | 3.2E-03 | phosphomethylpyrimidine synthase (ThiC) |

## Conclusions

Our work has shown that 15 transcription factors control the second tier of regulation in the quorum-sensing LuxR regulon (Fig. 5). Among these, several were previously known to control important cellular behaviors, such as type III secretion and osmotic stress response. The experiments in this study have expanded our knowledge of the LuxR-regulated genes in the quorum-sensing pathway of *V. harveyi* to include other pathways and metabolic systems, including methionine biosynthesis, thiamine biosynthesis, and acetate utilization. The second-tier of regulation controls at least 75 genes, though we did not examine the regulon of every gene (Fig. 5).

**Figure 5.**
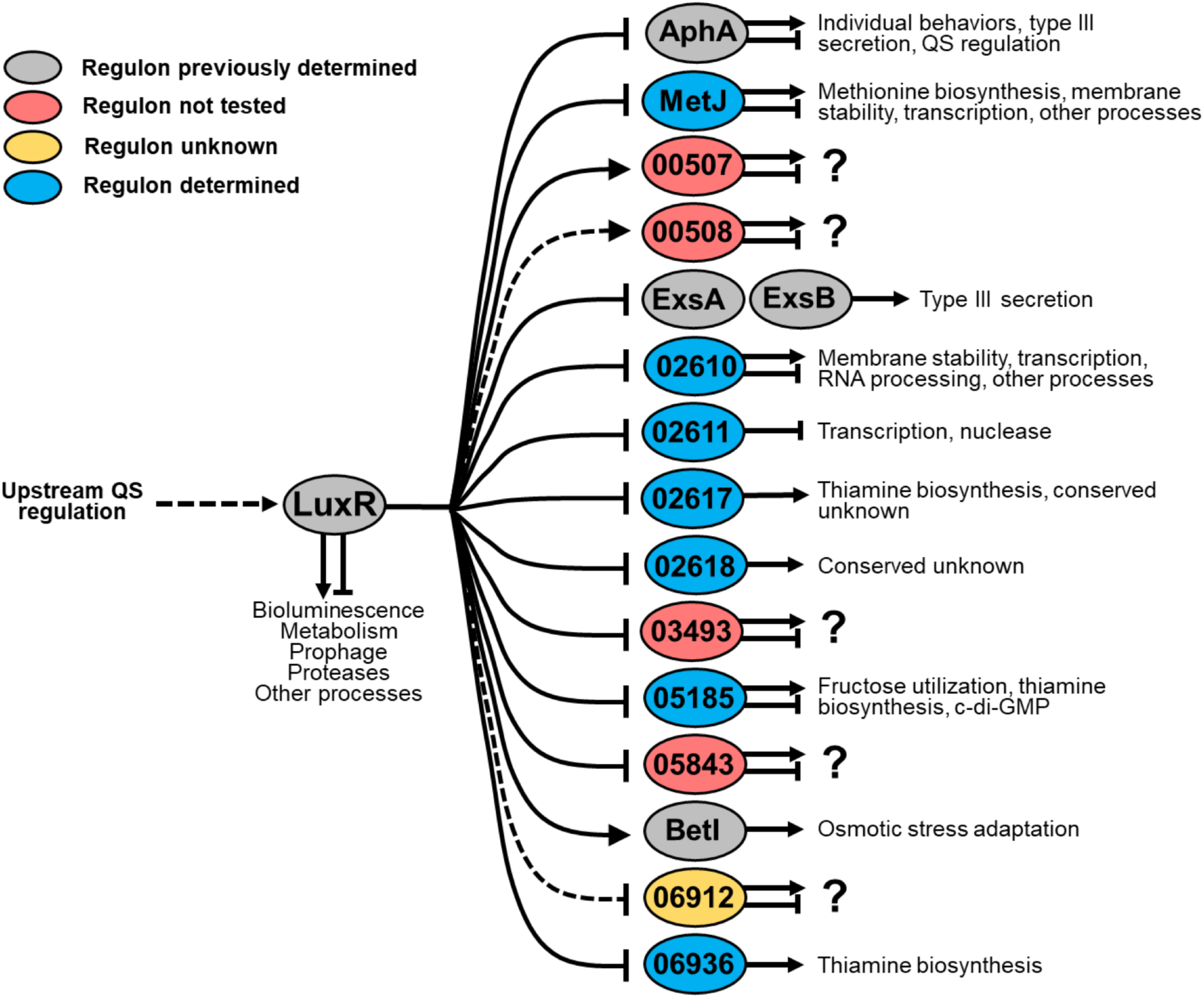
The hierarchy of LuxR regulation in *V. harveyi*. All LuxR-regulated transcription factors and their associated regulons are shown. Transcription factors are colored based on their respective regulons; gray indicates the regulon was determined previously (3, 11, 12, 13), red indicates the regulon was not tested in this study, yellow indicates the transcription factor did not regulate genes under our tested conditions, and blue indicates a regulon that was determined in this study. Arrows indicate positive regulation while blunt-end arrows indicate repression. Lines emanating from LuxR are solid to denote direct regulation or dotted to denote indirect regulation. Direct/indirect regulation of the second-tier regulons is not inferred in the model.

Some transcription factors regulated by LuxR have important connections to the quorum-sensing circuit in vibrios, including AphA in *V. harveyi* and VqsA in *Vibrio alginolyticus*. These regulators both feed back on the quorum-sensing system to regulate other components: 1) AphA represses the Qrr sRNAs and *luxR*, and the Qrrs activate *aphA*, and 2) VqsA activates *luxR* and represses *aphA* (3, 64). Of note, *V. harveyi* does not appear to encode a VqsA homolog; the closest conserved gene shares only 36% amino acid identity and is not regulated by LuxR (11, 12). Among the 8 transcription factors that we investigated, we did not observe evidence of feedback regulation onto any of the other quorum-sensing components (*e.g.*, *aphA* or the Qrrs). The experimental setup precluded us from evaluating regulation of *luxR* because the RNA-seq was performed in the Δ*luxR* background. However, other studies of Qrr targets did not reveal any of these transcription factor genes as regulated by the Qrrs (65). Thus, it is possible that there is not any regulatory connection of these transcription factors to the quorum-sensing circuit beyond LuxR-directed regulation.

Instead of feeding back onto the QS system, our data demonstrates that these LuxR-regulated transcription factors are involved in modulating several cellular processes. Among the genes that make up the second-tier of the LuxR regulatory cascade, many play a role in metabolism. This finding is not surprising when considering nutrient availability as cells reach quorum; at HCD, the nutrient profile available to the community changes and thus cells reconfigure metabolic programs, via LuxR, to respond accordingly. This is evidenced by the observation that acetate utilization genes are indirectly upregulated by LuxR. Acetate accumulation has been reported in *V. cholerae* cultures and HapR (its LuxR homologue) is a necessary component for mitigating the toxic effects of this organic acid (34). In addition to acetate utilization, our data has uncovered a clear connection between LuxR and methionine biosynthesis. LuxR may be responsible for maintaining methionine homeostasis as the cell transitions into a low-nutrient status. The discoveries made in this study contribute evidence to an evolving model of QS: amidst its many of responsibilities, reconfiguration of global cellular metabolism is yet another fundamental behavior that is controlled by LuxR and QS in *V. harveyi*.

## Materials and Methods

### Bacterial strains and methods

The strain of *V. harveyi* used is BB120 (ATCC BAA-1116), which was reclassified as *Vibrio campbellii* (66). For consistency with the literature, we refer to this strain as *V. harveyi* throughout this manuscript. *E. coli* strain S17-1*λpir* was grown at 30 °C with shaking (250 rpm) in Lysogeny Broth (LB) medium. *V. harveyi* and all mutant derivatives were grown at 30 °C with shaking (250 rpm) in Luria-Murine medium (LB medium with 2% total NaCl). Antibiotics were used at the following concentrations: 10 µg/mL chloramphenicol, 10 µg/mL tetracycline, 50 µg/mL polymyxin B, and 40 or 100 µg/mL kanamycin (*E. coli* or *V. harveyi*, respectively). Plasmid DNA was transformed into electrocompetent *E. coli* S17-1*λpir* cells using 0.2 cm electroporation cuvettes (USA Scientific). Transformed *E. coli* S17-1*λpir* were used to transfer plasmid DNA into *V. harveyi* via conjugation on LB plates at 30 °C overnight. Exconjugants were selected for using polymyxin B and other antibiotics described above on LM plates grown at 30 °C overnight. All strains used in this study are listed in Supplemental Table S1.

### Molecular methods

PCR was performed using Phusion HF polymerase (New England Biolabs). All restriction enzymes and T4 polynucleotide kinase were purchased from New England Biolabs and used according to the manufacturer’s protocol. All oligonucleotides were ordered from Integrated DNA Technologies. Oligonucleotides used for EMSAs and qPCR are listed in Supplemental Table S3. Plasmids were constructed via isothermal DNA assembly (IDA; all enzymes and dNTPs from New England Biolabs). PCR products and plasmids were cleaned up using Qiagen kits and sequenced using Eurofins Genomics. All plasmids used in this study are listed in Supplemental Table S2. DNA samples were resolved using 1% agarose and 1x TBE buffer.

### Mutant construction

*V. harveyi* mutants were constructed using the protocol as described (12). Briefly, 1000 bp upstream and downstream of the target gene were cloned into the suicide vector pRE112 flanking a chloramphenicol (CM) resistance marker and a *sacB* counterselectable marker. pRE112 contains oriR_R6Kγ_, which allows for replication in *E. coli* S17-1*λpir* but not *V. harveyi*. Plasmids were conjugated into *V. harveyi* from *E. coli* and exconjugants selected on CM-containing LM plates. Once integrated, plasmid excision was selected for by streaking cells on LM plates containing 20% sucrose. Isolated colonies were screened using PCR to identify cells in which plasmid excision yielded a successful gene deletion. All mutants were confirmed with sequencing.

### RNA isolation and qRT-PCR

For RNA-seq and qRT-PCR experiments, cells were grown to OD_600_ = 1.0 and 5 mL of culture were harvested, pelleted, and frozen on liquid N_2_. RNA isolation was performed as described in previous studies using a Trizol/chloroform extraction method (3, 12). DNA in samples was digested and removed using DNaseI (Ambion RNase-free kit). The RNA sample was purified using an RNeasy purification kit (Qiagen). The Sensi-FAST SYBR Hi-ROX One-Step Kit (Bioline) was used for quantitative real time PCR (qRT-PCR). Reactions were set up according to the manufacturer’s protocol and 50 ng of purified RNA from cultures was added to each reaction in a 96-well plate (Applied Biosystems). qRT-PCR was performed using a StepOnePlus Real-Time PCR system (Applied Biosystems) using thermal cycling conditions according to the SensiFAST protocol (Bioline). The *hfq* gene was used as an internal standard for normalization as its expression is constitutive (and unaffected by LuxR) in *V. harveyi*. To analyze relative fold-changes in RNA levels, the standard curve method was used on data acquired from three independent biological replicates.

### RNA sequencing

RNA-seq analysis was performed at the Center for Genomics and Bioinformatics at Indiana University using RNA that was purified as described above. RNA-seq was performed as described previously (12). Sequence data were deposited in the National Center for Biotechnology Information Gene Expression Omnibus database (NCBI GEO: GSE144017).

### Electrophoretic mobility shift assays (EMSAs)

DNA substrates for EMSA experiments were obtained via PCR. PCR products were cleaned with a purification kit (Qiagen) and the 5’ ends of the DNA (25 nM) were radiolabeled using T4 polynucleotide kinase (PNK, New England Biolabs). Excess, unincorporated ATP [γ-^32^P] was removed using G-50 spin columns (GE Life Technologies) following the manufacturer’s protocol. DNA binding reactions were performed using 1 nM 5’-radiolabelled DNA in LuxR binding buffer (10 mM HEPES pH 7.5, 100 mM KCl, 2 mM DTT, 200 µM EDTA). All reactions included 100 µg/mL bovine serum albumin (BSA) and 10 ng/µL poly(dI-dC). Reactions were incubated at 30 °C for 30 min and then run on 6% TGE non-denaturing gel. Gels were dried at 80 °C for 20 minutes and exposed to a phosphor screen overnight. Phosphor screens were visualized using a Typhoon 9210 (Amersham Biosciences).

## Acknowledgments

We thank Chelsea Simpson and Victoria Lydick for excellent technical assistance. We also thank Douglas Rusch, Jun Liu, and Dave Miller at the Indiana University Center for Bioinformatics and Genomics for assistance with RNA-seq sample processing and analyses. We thank Sofia Quinodoz for her insights and discussion of the data.

